# A large close relative of *C. elegans* is slow-developing but not long-lived

**DOI:** 10.1101/426254

**Authors:** Gavin C. Woodruff, Erik Johnson, Patrick C. Phillips

## Abstract

**Background:** Variation in body size is thought to be a major driver of a wide variety of ecological and evolutionary patterns, including changes in development, reproduction, and longevity. *Caenorhabditis inopinata* is a recently-discovered fig-associated nematode that is unusually large relative to other members of the genus, including the closely related model system *C. elegans*. Here we test whether the dramatic increase in body size has led to correlated changes in key life history and developmental parameters within this species.

**Results:** Using four developmental milestones, *C. inopinata* was found to have a slower rate of development than *C. elegans* across a range of temperatures. Despite this, *C. inopinata* did not reveal any differences in adult lifespan from *C. elegans* after accounting for differences in developmental timing and reproductive mode. *C. inopinata* fecundity was generally lower than that of *C. elegans*, but fitness improved under continuous-mating, consistent with sperm-limitation under gonochoristic (male/female) reproduction. *C. inopinata* also revealed greater fecundity and viability at higher temperatures.

**Conclusion:** Consistent with observations in other ectotherms, slower growth in *C. inopinata* indicates a potential trade-off between body size and developmental timing, whereas its unchanged lifespan suggests that longevity is largely uncoupled from its increase in body size. Additionally, temperature-dependent patterns of fitness in *C. inopinata* are consistent with its geographic origins in subtropical Okinawa. Overall, these results underscore the extent to which changes in ecological context and body size can shape life history traits.

## Background

Trade-offs dominate life history evolution. Organisms have access to limited energy resources, and these must be allocated in a balance between self-maintenance and reproductive output. In keeping with the expectation that different distributions of life history traits (such as age of maturity, reproductive duration, and age-specific fecundity, among others) should be sensitive to different distributions of selective pressures on those traits, a huge diversity of patterns among life history traits has emerged across the broad scope of animal diversity [1–5]. As a consequence, many organisms exhibit well-documented correlations among traits such as fecundity and survival [6–8], fecundity and developmental rate [1, 9–11], and reproductive quantity and quality [12, 13].

Body size is a particularly potent component of life history syndromes. Body size is usually correlated with a multitude of fitness-related traits including developmental rate, offspring number, offspring size, gamete size, and lifespan [14–17]. Body size is also known to covary with physiological traits, such as metabolic rate, thought to underlie trade-offs among life history traits [15, 17]. These factors in turn generate allometric relationships that appear to explain scale-based trends for a wide variety of traits across many orders of magnitude [15]. Indeed, body size appears to be a central component of broad macroevolutionary trends among lineages over geological timescales [18]. But which is cause and which is effect? To what extent does change in body size due to selection on body size per se lead to collected changes in such a wide array of life history traits and to what extent does body size change because of selection acting directly on these traits?

Life history theory suggests that selection for increased body size can be balanced against the benefits of faster reproduction and the costs of lower offspring viability and lower initial fecundity [1], weighed against a backdrop of differential allocation of physiological and metabolic resources to each of these processes and to growth itself [17, 19]. At the same time, selection on body size itself must be mediated via environmental factors such as resource availability and/or predation [20]. Although these various causes are not mutually exclusive and likely overlap, the proximate and ultimate causes of changes in body size change—particularly the relationship between these two—remain largely unresolved.

The nematode *Caenorhabditis elegans* has for decades been an important model for genetics, development, and biology in general [21]. However, the degree and extent of trade-offs between body size and other life history traits in systems like *C. elegans* remain largely unknown and/or have generated somewhat ambiguous or contradictory results [22–30]. Further, because nearly all known members of this genus share a common natural ecological niche of rotting plant material [31], it has not been possible to use a comparative approach to investigate how change in ecological circumstances might drive changes in the relationship between body size and life history [19]. Here, we address this question by taking advantage of a highly phenotypically and ecologically divergent close relative of *C. elegans*: the recently discovered fig-associated nematode *C. inopinata*.

*C. inopinata* (formerly known as *C.* sp. 34) is remarkable in that it displays tremendous ecological and phenotypic differences compared to its close relatives [32, 33]. Compared to other *Caenorhabditis*, *C. inopinata* is huge: it can grow to be nearly twice as long as other members in the genus [32, 33]. *C. inopinata* also develops nearly twice as slowly, has sperm three times the size, and embryos 20% longer than *C. elegans* [33]. Furthermore, in contrast to the rotting-plant material ecological niche of *C. elegans* and other *Caenorhabditis* species [34], it thrives in the fresh, intact Okinawan figs of *Ficus septica* [32, 33, 35]. *C. inopinata* thus appears to have experienced a radically different selective environment that has led to its highly divergent suite of life history traits. And, as *C. inopinata* is much larger in size and develops much more slowly than its close relatives, it can therefore be used as a natural system to test the predictions of life history theory using a comparative approach. Here, we do just this by describing the developmental timing, lifespan, fecundity, and viability of *C. inopinata* and *C. elegans* at multiple temperatures.

## Results

### C. inopinata *develops more slowly yet does not differ from* C. elegans *in lifespan and reproductive duration*

Initial measures of developmental rate revealed that *C. inopinata* develops about twice as slowly as *C. elegans* [33]. To provide a more complete picture of the timing of development in this species, the occurrence of four different developmental milestones (time of hatching, onset of the L4 stage, onset of adulthood, and the onset of reproduction) was ascertained at four different temperatures (15°C, 20°C, 25°C, and 30°C) among synchronized populations of *C. elegans* and
*C. inopinata*. Unsurprisingly, all species grew faster as the temperature increased (Figure 1; Table S1). Yet in conditions where both species grew reliably, *C. inopinata* was slower to reach all developmental milestones than *C. elegans* (Figure 1; Table S1). Indeed, at the typical rearing temperature of *C. elegans* (20°C), the median time of reproductive onset was 2.7 days in *C. elegans*, whereas it was 6.7 days in *C. inopinata* (GLM LRT chi-square=4861.4, df=2, p<0.0001). To reach a developmental rate that approaches that of *C. elegans* at 20°C, *C. inopinata* must be reared at a temperature that is ten degrees higher (Figure 1b; Table S1) where it exhibits reduced fecundity (Figure 4a) and where *C. elegans* N2 is inviable (Figure 5). Overall, then, *C. inopinata* has slower relative growth regardless of temperature.

**Figure 1.**
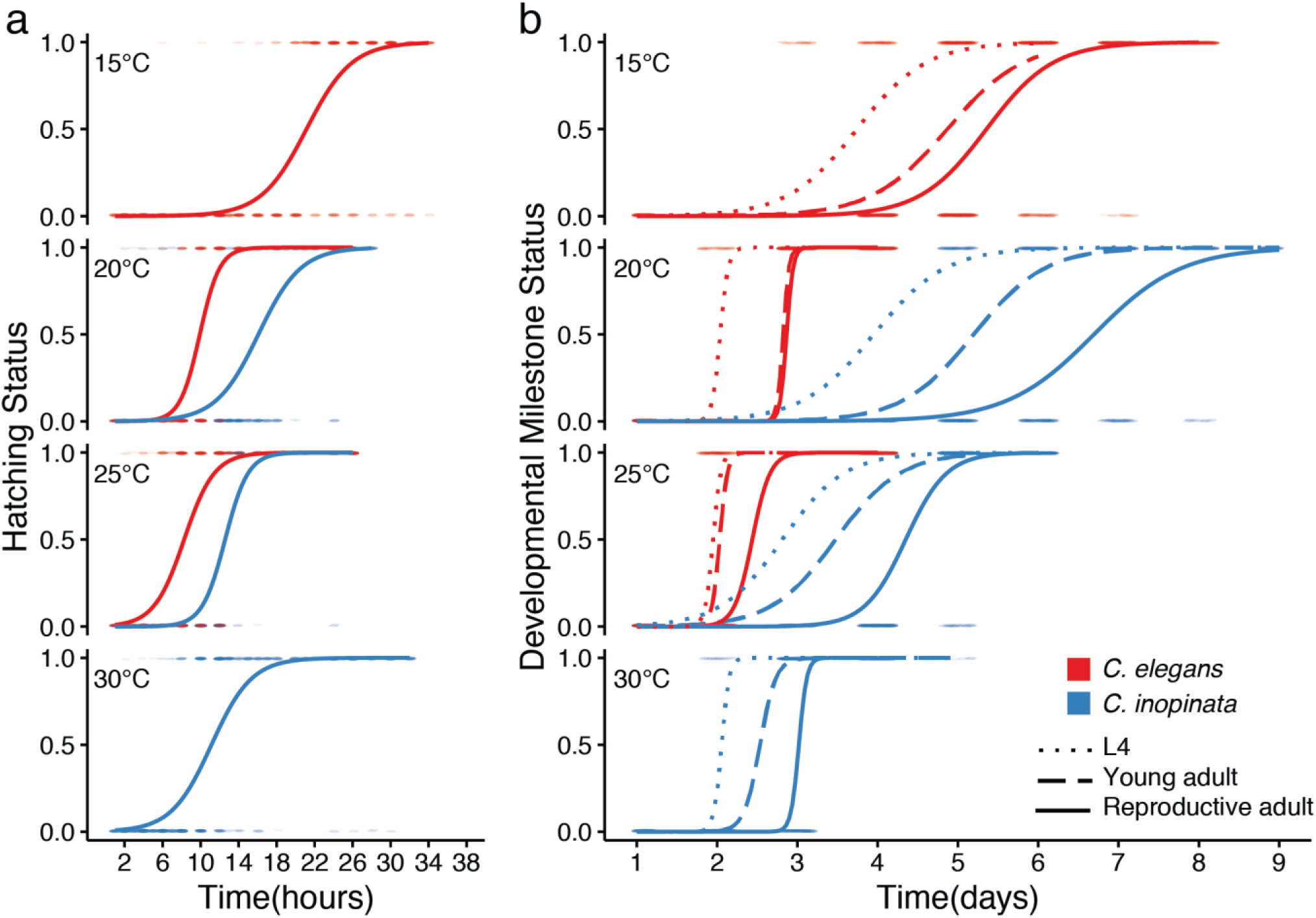
*C. inopinata* develops more slowly than *C. elegans*. The y-axis represents the status of having attained a given developmental milestone; 0 = yet to reach milestone, 1 = has reached milestone. a) Hatching; b) L4, young adulthood, and the onset of reproduction. *C. elegans (fog*-*2)* was used for the embryogenesis milestone to account for the delay caused by obligate outcrossing in *C. inopinata*. *C. elegans* N2 is inviable at 30°C, and *C. inopinata* milestones were not measured at 15°C due to its low fitness at this temperature. N worms = 186-960. GLM LRT chi-square p<0.0001 for every *C. elegans* and *C. inopinata* comparison.

As slow developing, large animals tend to be longer-lived [1], we were curious if *C. inopinata* also exhibits prolonged longevity. To address this, we applied previously established methods of lifespan measurement in nematodes [36] to *C. inopinata*. As a point of comparison, we also measured *C. elegans* N2 and *C. elegans* (*fog*-*2*) lifespans. As lifespan often trades-off with reproductive output [37, 38], we used virgin *C. elegans (fog*-*2)* pseudo-females (which do not generate self-sperm and are self-sterile as a consequence [39]) to control for differences in reproductive mode. *C. inopinata* females were longer-lived than wild-type *C. elegans* hermaphrodites at 25°C, with a median total lifespan that was four days higher (20 and 16, respectively; Cox proportional hazards linear model comparison, Z-value=4.99, p<0.0001 Figure 2a; Figure S1). However, *C. inopinata* females were only marginally longer lived than *C. elegans (fog*-*2)* pseudo-females (19 days, Cox proportional hazards linear model comparison, Z-value=2.29, p=0.053). Furthermore, no differences in adult lifespan (which takes into account the differences in developmental timing between *C. elegans* and *C. inopinata*) were detected between *C. inopinata* females (median adult lifespan of 16 days) and *C. elegans (fog*-*2)* pseudo-females (median adult lifespan of 17 days; Cox proportional hazards linear model comparison, Z-value=0.74, p=0.73; Figure 2b; Figure S2). Thus, despite its large size and slow development, *C. inopinata* adults are not longer-lived than *C. elegans* after accounting for differences in reproductive mode and developmental timing.

**Figure 2.**
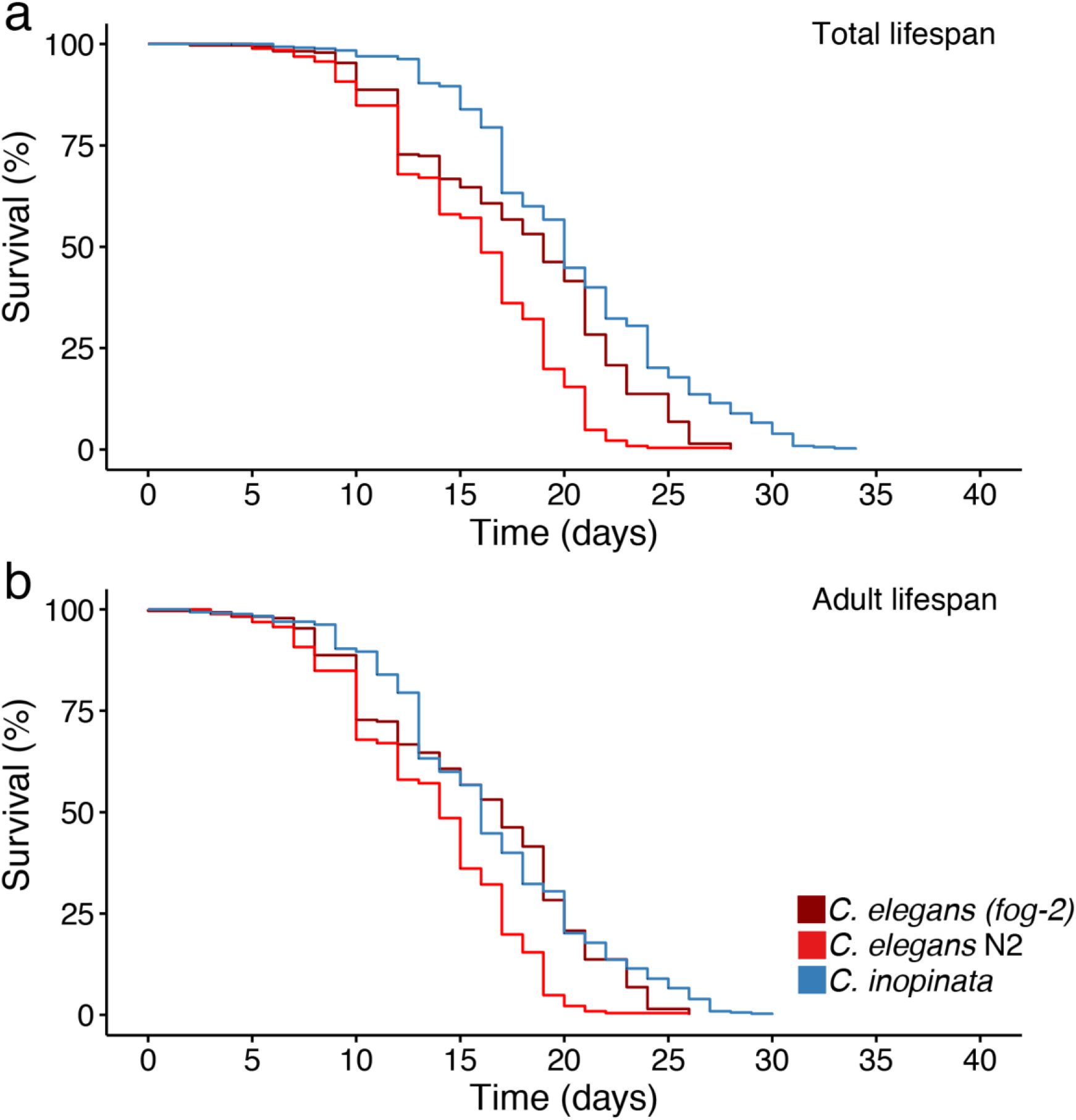
*C. inopinata* is not longer-lived than *C. elegans* at 25°C after taking reproductive mode and developmental timing into account. (a) Total lifespan models. Here, Day = 0 represents the day embryos were laid. (b) Adult lifespan models. Here, Day = 0 is the approximate first day of adulthood, taken as the total lifespan minus two (*C. elegans*) or four (*C. inopinata*) days. Wild-type *C. elegans* N2 exhibits both shorter total and adult median lifespan than *C. inopinata*. Conversely, *C. inopinata* females have a marginally higher median total lifespan than non-selfing *C. elegans (fog*-*2)* mutant females, and no difference in *C. inopinata* and *C. elegans (fog*-*2)* adult lifespan was detected (Cox proportional hazards linear model comparison, Z-value=0.74, p=0.73). N worms=263 (*C. elegans* N2), 281 (*C. elegans (fog*-*2)*), 444 (*C. inopinata*).

The duration of reproduction is also expected to trade-off with growth rate and body size [1, 2], with large, slow-developing animals tending to have longer reproductive periods [9–11]. To see if this also holds for *C. inopinata*, daily measures of fecundity were made with individual *C. elegans* (*fog*-*2*) pseudo-females and *C. inopinata* females under conditions of continuous mating throughout their lifetimes (Figure 3). Although one individual *C. inopinata* female had a reproductive duration of twelve days, for the most part, both species lay almost all of their embryos in the first four days of adulthood (Figure 3b). Indeed, under continuous mating conditions at 25°C, no differences in brood fraction per day could be detected between *C. inopinata* and *C. elegans* with the exception of day eight of adulthood (Wilcoxon rank sum test, W=528, p=0.041). Thus, like lifespan, duration of reproduction is not extended in *C. inopinata*.

**Figure 3.**
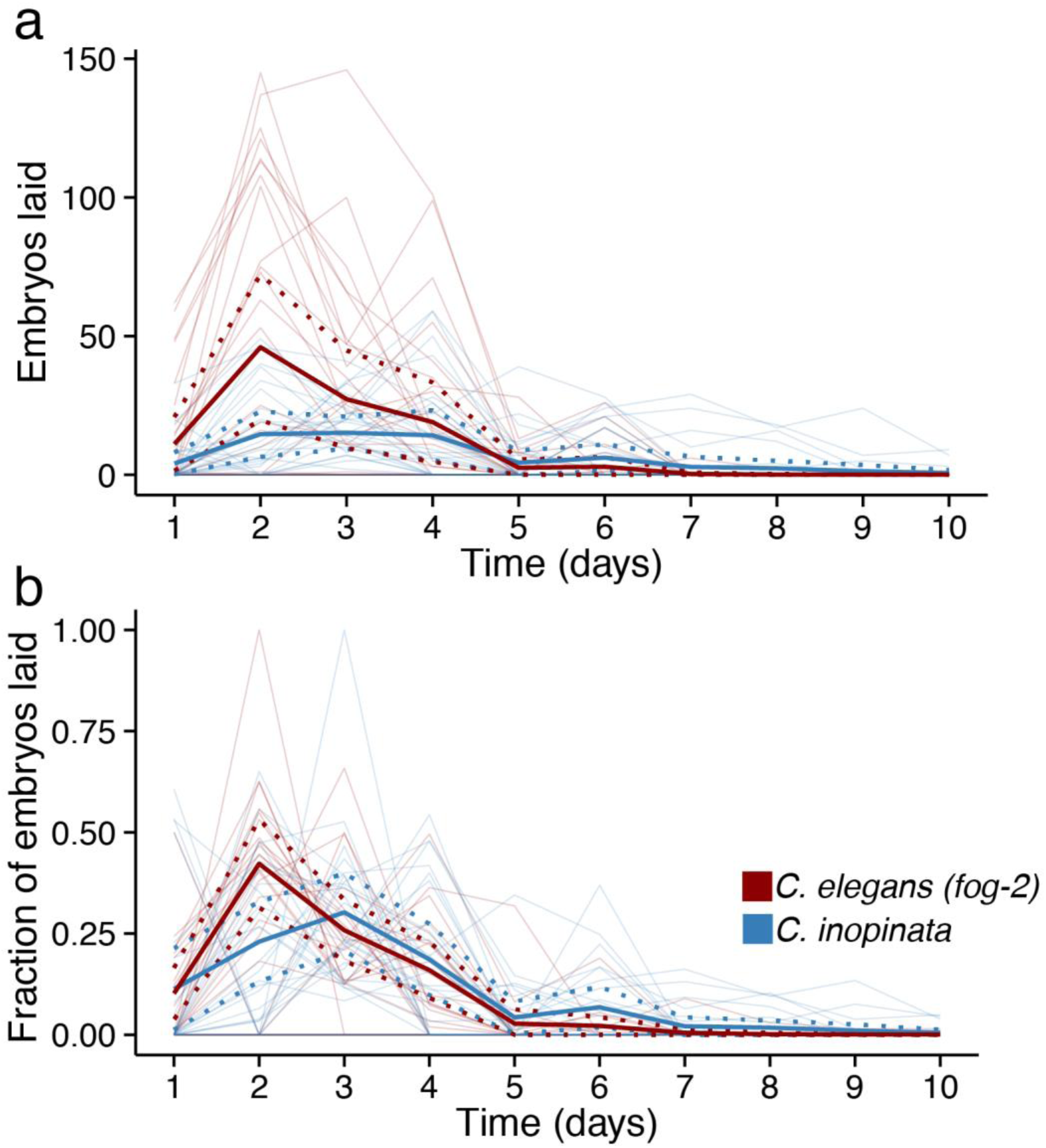
*C. inopinata* has a reproductive duration comparable to *C. elegans*. (a) Numberof embryos laid per day. (b) Fraction of lifetime brood laid per day. Bold lines represent averages, and dotted bold lines represent ±1 SDM. Thin lines represent individual worms. The *C. elegans (fog*-*2)* and *C. inopinata* day two and three brood fractions are not statistically different (Wilcoxon rank sum test W=389 p=0.36 and W=553 p=0.13, respectively). N parental females=30 for both species. All observations taken at 25°C.

### C. inopinata *is sperm-limited and reveals higher fitness at higher temperatures*

Brood size also tends to covary with both body size and developmental rate [1, 2], and so fecundity was measured at four different temperatures in *C. inopinata* and *C. elegans (fog*-*2)* to address if similar patterns hold in this group (Figure 4). In conditions in which females were mated with males for just one night, *C. inopinata* generally displayed far smaller brood sizes than *C. elegans (fog*-*2)*, with the exception that *C. elegans (fog*-*2)* is infertile at 30°C (Figure 4a). However, as the male/female species *C. remanei* is known to generate more progeny when constantly exposed to males [40, 41], we suspected that *C. inopinata* might also be sperm-limited. Indeed, under continuous mating conditions, there is no detectable difference in brood size between *C. inopinata* and *C. elegans (fog*-*2)* (median brood size of 58 and 76, respectively; Wilcoxon rank sum test, W=484 p=0.62; Figure 4b). However, male mating performance tends to degrade in selfing species [42], so we also compared the fraction of successful crosses between *C. elegans* and *C. inopinata* (Figure S3). In continuous mating conditions, the fraction of failed crosses was higher in *C. elegans* (0.33, N=30 crosses) than in *C. inopinata* (0.17, N=30 crosses), although this difference was not statistically significant (Fisher’s Exact Test odds ratio=2.46, p=0.23). After removing animals that failed to produce progeny, *C. elegans (fog*-*2)* yielded a median brood size that is over twice as large as that of *C. inopinata* in continuous mating conditions (145 and 65, respectively; Wilcoxon rank sum test, W=359, p=0.013; Figure S4). Thus *C. inopinata* requires constant access to mates in order to maximize its reproductive output, consistent with its gonochoristic mode of reproduction.

**Figure 4.**
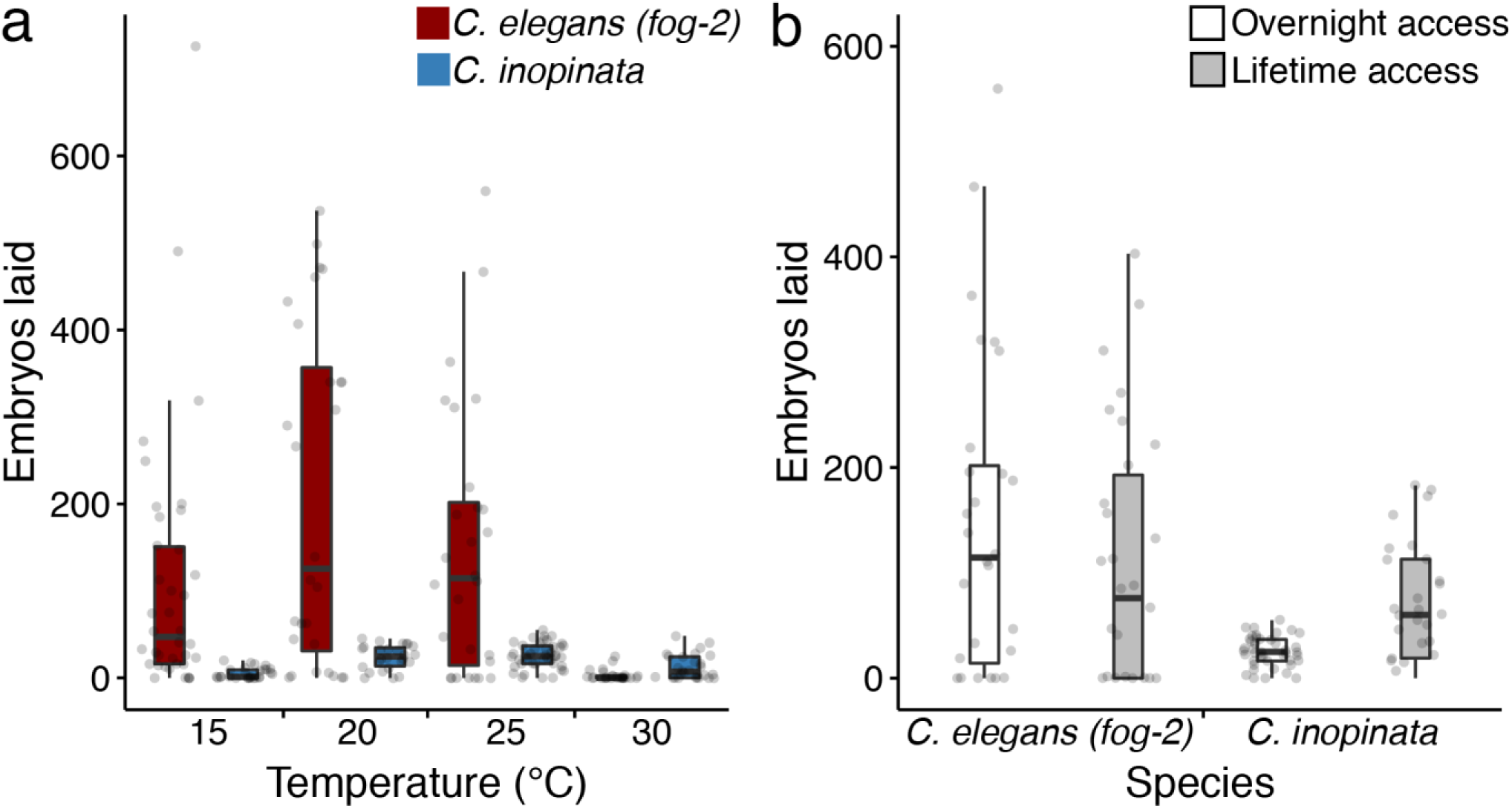
*C. inopinata* is sperm-limited. (a) Number of embryos laid in single overnight mating conditions at various temperatures. (b) Number of embryos laid in continuous mating or single overnight mating conditions at 25°C. The “one overnight mating” data in panel (b) is the same from those at 25°C in panel (a). *C. inopinata* has smaller broods than *C. elegans (fog*-*2)* in every condition except 30°C (Wilcoxon rank sum test p<0.0001 for 15 and 20°C; W=349, p=0.004 for 25°C; W=575, p=0.002 for 30°C). However, there is no detectable difference in *C. elegans (fog*-*2)* and *C. inopinata* brood sizes under continuous mating conditions (Wilcoxon rank sum test, W=484, p=0.62). N parental females=26-42.

When examining the relationship between developmental rate and fecundity, the intrinsic rate of increase (*r*) is likely a better measure of fitness than total fecundity (*R*_0_) [1, 43]. Under this approach, fitness is a function of age-specific fecundity and viability, and the age of first reproduction can highly influence the population growth rate [1]. So although *C. inopinata* and *C. elegans* have comparable brood sizes under continuous mating conditions, they likely differ in fitness because of their different developmental rates. Indeed, despite their comparable brood sizes, *C. elegans* has a rate of increase (*r*=1.54, 95% CI=1.26-1.72) that is over twice as high as *C. inopinata* (*r*=0.66, 95% CI=0.54-0.74). This difference in fitness is even greater in mating conditions with just overnight access to males (*C. elegans r*=2.09, 95% CI=1.88-2.24; *C. inopinata r*=0.63, 95% CI=0.55-0.69). Thus continuous access to males is not sufficient to overcome the detriment to fitness due to slow development in *C. inopinata*.

In keeping with the other life-history measures, *C. elegans* was more viable at lower temperatures and *C. inopinata* more viable at higher temperatures during early development (Figure 5). Overall, however, *C. inopinata* displayed consistently lower embryo-to-adult viability than *C. elegans* at 15°C, 20°C, and 25°C (Wilcoxon rank sum test p<0.001 in all comparisons; Figure 5). No detectable differences in *C. inopinata* viability were found between 20°C, 25°C, and 30°C (median viability of 0.84, 0.79, and 0.88, respectively; Wilcoxon rank sum test W=50 p=0.060, W=70 p=0.62; Figure 5), but *C. inopinata* is less viable at 15°C (median viability of 0.63; Wilcoxon rank sum test p≤0.030 for all comparisons). As *C. inopinata* fecundity is also higher at warmer temperatures (Figure 4a), these temperature-specific fitness patterns are consistent with its subtropical natural context of fresh Okinawan *Ficus septica* figs.

**Figure 5.**
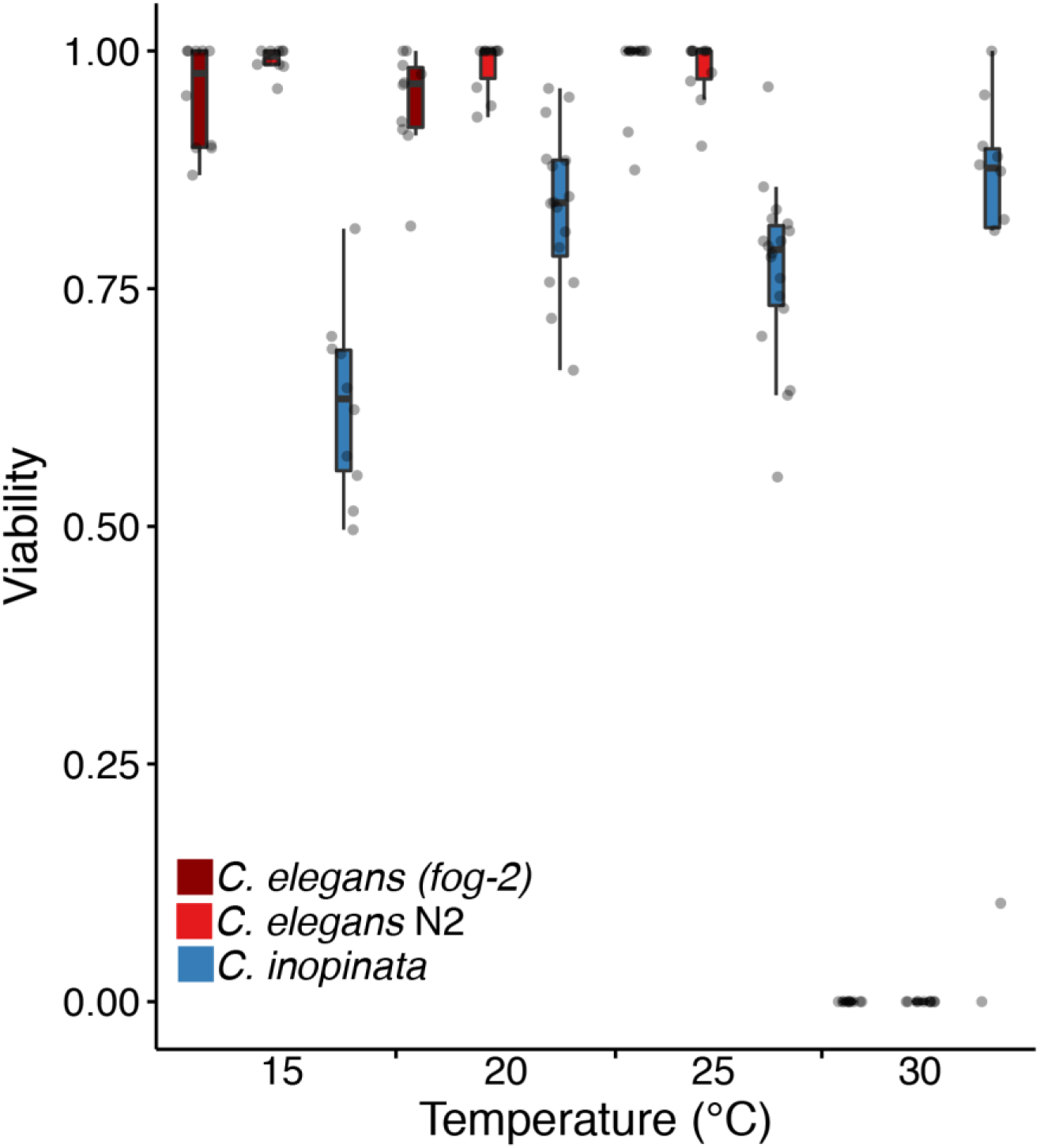
*C. inopinata* has a lower viability than *C. elegans*. Embryo-to-adult viability at four temperatures. *C. elegans* reveals higher viability in all conditions except 30°C regardless of reproductive mode. N observed plates of worms=10-18 per strain per condition; N embryos per plate=5-237.

## Discussion

Possibly because it is both obvious and easy to measure, body size variation has been studied extensively for centuries. The range in body size across the tree of life is so immense as to demand explanation (21 orders of magnitude [16, 44]), and this incredible diversity has spawned a vast and rich literature attempting to comprehend its origins and maintenance. One major conclusion from this research program is that body size is correlated with nearly every trait, such that long-established relationships between body size and growth, reproduction, and lifespan underscore a prominent role for body size in the evolution of life histories [14, 15, 44]. Here, we found that an exceptionally large close relative of *C. elegans* exhibits slow growth and low fecundity across a range of temperatures yet is not long lived. Together with the extensive *C. elegans* literature and the foundations of life history theory, these observations can inform our understanding of the causes and consequences of large-scale changes in body size.

### Developmental timing

It makes intuitive sense that larger organisms should develop more slowly. Being more massive, presumably more cell divisions and/or biosynthetic reactions must take place for their construction and it therefore follows that their development should take longer than smaller organisms. And this intuition bears out across vast phylogenetic distances: from bacteria to sequoias, body size covaries with generation time [44]. Here, we found that in all temperatures, *C. inopinata* grows nearly twice as slowly as *C. elegans*, consistent with previous observations (Fig. 1; [32, 33]). Indeed, *C. inopinata* needs to be grown at 30°C to approach a rate of development comparable to that of *C. elegans* when grown at 20°C. Thus, the observation that this very large species also develops much more slowly than its close relatives is in line with decades of allometric studies. Further, as cell size is coordinated with cell division decisions in multiple organisms [45, 46], body size change could occur even in the absence of cell number change through the modification of cell cycle timing. This may explain the case of *C. inopinata*, as previous observations observed no change in cell number despite its large size and slow development [33].

However, there are reasons to suspect slow development may not underlie large body size in this case. It has been argued that the allometric trends observed in birds and mammals cannot be easily extended to poikilotherms because of difficulties in comparing physiological time due to rapid change in metabolic rates [16]. More notable is the common observation that developmental timing can be decoupled from body size in *C. elegans*. Most mutations in *C. elegans* that extend body length do not also slow the rate of growth: only 29% of the genes in the *C. elegans* genome known to control body length also promote slower development (Figure S5). Furthermore, experimental evolution and mutation accumulation studies in *C. elegans* and *C. briggsae* have not generally reported correlated changes in body size and developmental timing [23, 25, 26, 47]. Thus, it appears that body size and rate of growth need not be strongly coupled in *Caenorhabditis* and that the relationship between these traits observed in *C. inopinata* may not necessarily be causative.

Instead, the slow growth of *C. inopinata* may be better understood with respect to its natural ecological context. *C. inopinata* is associated with fresh figs and their pollinating wasps [35], whereas their close relatives tend to proliferate on rotting plant material [34]. And as *C. inopinata* animals disperse to new figs via pollinating wasps [35], their life cycle is necessarily closely tied to patterns of wasp development and emergence. Figs generally take weeks to develop [48], and although it is unclear how many generations of worms occur within a single fig, it is reasonable to suspect that the extreme divergence in developmental rate is connected to its novel natural context. This is consistent with correlations among *Ceratosolen* fig wasp and *C. inopinata* developmental stages that have been found in previous field studies [35]. Future longitudinal field studies of single fig trees at finer temporal resolution will be required to determine the relative paces of fig, fig wasp, and nematode development in nature and to test hypotheses regarding the ecological drivers of heterochrony.

### Reproduction

The relationship between body size and reproduction varies both within and between taxa. In birds and mammals, larger species tend to have lower fecundities than smaller species [15]. Conversely, body size appears to be positively correlated with fecundity in insects [49] and nematodes [50]. *C. inopinata* was generally found to have lower brood sizes than *C. elegans* across a range of temperatures (Fig. 4a), although continuous mating greatly improves fecundity in *C. inopinata* (Fig. 4b). The relatively low fecundity of *C. inopinata* is then incongruent with patterns of fecundity and body size that have been generally observed in nematodes. *C. inopinata*’s gonochoristic mode of development cannot explain its low brood size, as multiple male/female species of *Caenorhabditis* have been reported to have higher brood sizes [40, 41, 51–54]. However, the sperm-limited fecundity of *C. inopinata* (Fig. 4b) is consistent with previous observations with the gonochoristic *C. remanei* [40, 41]. It is possible that the evolution of extreme body size in the case of *C. inopinata* reveals a trade-off with reproductive output, wherein resources usually allocated to progeny have instead been shifted to increase self-maintenance and growth. Yet most genes known to regulate body length in the *C. elegans* genome do not have a pleiotropic role in brood size (only 28% do; Figure S5). This is also consistent with experimental evolution studies in *Caenorhabditis* [23], wherein fecundity and body size do not necessarily trade-off. So again, the precise causal relationship here bears further study.

A particularly interesting avenue to pursue is based on the observation that wild bacteria associated with *Caenorhabditis* can have both positive or negative influences on fecundity and growth [55, 56] and that different species of *Caenorhabditis* are associated with different microbes in nature [55]. Thus the nutritional environment can have a profound effect on fitness. The natural microbial food of *C. inopinata* is currently unknown. As *C. inopinata* exhibits reduced gonads in laboratory culture [33], it may be experiencing nutritional deficiencies. The reduced fecundity of *C. inopinata* may then reflect a plastic response to an adverse environment as opposed to a trade-off with increased body size. The potential influence of natural microbial associates of *Ficus septica* figs on *C. inopinata* fitness affords an exciting opportunity for future research.

### Lifespan

Lifespan is often positively correlated with body size, and from an allometric perspective is usually thought to be regulated by variation in developmental and metabolic rates [15, 17]. And although the age of maturity is sensitive to selection under a range of trait distributions in life history theory [1], from an evolutionary perspective it is thought that late-life traits are generally not subject to selection as its strength falls to zero once reproduction ends [3]. Despite its large size and slow development, *C. inopinata* was found to have only a marginally longer lifespan than *C. elegans* (Fig. 2). And, when differences in developmental timing and reproductive mode are taken into account, *C. inopinata* adult lifespan is not significantly different from that of *C. elegans* (Fig. 2b). The lack of lifespan change in this system is consistent with the view that lifespan is under weak selection, as *C. inopinata* has experienced dramatic change in many other traits under its novel ecological context [32, 33, 35]. Indeed, most lifespan-extending mutations identified in *C. elegans* have not been associated with pleiotropic effects on body size (Figure S5). Similarly, experimental evolution studies in *C. elegans* show no correlated responses in lifespan upon artificial selection on early fecundity [30] and body size [23]. Additionally, no relationships between lifespan and fecundity have been found in mutation-accumulation lines [22] or among wild isolates [24]. These observations are inconsistent with the antagonistic pleiotropy explanation of aging, which posits that the greater fitness contribution of early life survival and reproduction leads to late life deterioration because of negative genetic correlations of these traits [57]. Rather, lifespan appears to be possibly largely uncoupled from fitness-related traits in this group, consistent with the unchanged longevity observed in *C. inopinata*. However, the nutritional caveats in this system noted in the above interpretation of observed patterns of fecundity also apply here. It is possible that *C. inopinata* will be longer-lived under different rearing conditions, and measurements of lifespan of *C. inopinata* raised on bacterial food originating from its natural context need to be performed.

### Temperature-dependent patterns of fitness-related traits in C. inopinata

Notably, *C. inopinata* was more fit at higher than lower temperatures (Fig. 4a, Fig. 5). Temperature-dependent plasticity of fitness-related traits varies both within and between species in *Caenorhabditis*, and these patterns often coincide with ecological context. Within *C. briggsae*, there are definable clades that are genetically structured by latitude [58, 59], and these wild isolates reveal temperature-dependent patterns of fecundity that are consistent with their geographical origin [60]. Additionally, the tropical species *C. nigoni* [51, 61] and *C. tropicalis* [62] have higher fitness at warmer temperatures. As *C. inopinata* has only been found in the subtropical islands of Okinawa [32, 33], its temperature-dependent patterns of fitness are consistent with these previous observations. And further, the temperatures where *C. inopinata* has shown the highest fitness here are comparable to natural *Ficus septica* fig temperatures measured in nature [35]. As a close relative of *C. elegans*, this species is well positioned for uncovering the genomic bases of temperature adaptation.

## Conclusions

Body size is a major driver of evolutionary change in multiple taxa, and changes in body size often co-occur with widespread change in life history traits. Here, we examined the life history traits of a large, ecologically-divergent close relative of *C. elegans*. We found that *C. inopinata* develops nearly twice as slowly as *C. elegans*, revealing a likely trade-off between growth and body size. Conversely, longevity does not evolve as part of correlated response to selection on body size in this system, consistent with previous studies and indicative of genetic decoupling of longevity from other life-history traits. Furthermore, patterns of fecundity in *C. inopinata* are also inconsistent with those expected of large nematodes. Hence change in body size alone cannot predict the evolution of whole suites of life history traits. Future studies that situate these systems within their natural ecological contexts will be needed to fully disentangle matters of cause and effect among the traits that constitute life history strategies. Taken together, these observations reveal that drastic change in ecological context and body size do not necessarily have an all-encompassing impact on life history syndromes.

## Methods

### Strains and maintenance

Animals were maintained on Nematode Growth Media (with 3.2% agar to discourage burrowing) supplemented with *Escherichia coli* strain OP50-1 for food. The *C. inopinata* wild isolate strain NKZ2 [33] was utilized for all observations in this report. *C. elegans* N2 and the obligate outcrossing *C. elegans fog*-*2(q71)* JK574 [39] mutant strain were also used for most comparisons. Notably, *C. elegans* is hermaphroditic, while *C. inopinata* is male/female or gonochoristic. This makes interspecific comparisons problematic. Thus the *fog*-*2(q71)* mutation, which prevents spermatogenesis only in hermaphrodites but promotes no obvious somatic defects in either sex [39], was used to control for differences in reproductive mode in various comparisons of life history traits.

### Developmental timing

The timing of four developmental milestones (hatching, L4 stage, adult stage/young adulthood, and the onset of reproduction/reproductive adulthood) was measured at four temperatures: 15°C, 20°C, 25°C, and 30°C. For synchronization, mid-stage embryos (blastula to 1.5 fold stage) were picked from plates cultured at 25°C to new plates and then shifted to the given rearing temperature. Plates were then monitored hourly (for hatching) and then daily (for L4, young adulthood, and reproductive adulthood) for the onset of developmental milestones. Male tail and female/hermaphrodite vulva morphologies were used to define L4 and young adult stages. The onset of reproduction was scored only among females and hermaphrodites by the presence of embryos in the uterus. Plates were assayed until the number of individuals at or older than a given milestone did not increase for two hours or days. Animals who failed to reach a given milestone were not used for subsequent analysis. For analysis, animals were plotted by their developmental status (“0” = yet to reach milestone; “1” = reached milestone) over time and logistic regression was used to estimate the median time to a given event via the “glm” function (using a binomial distribution) in the R statistical language. This models approach was used for hypothesis testing and for calculating 95% confidence intervals (see Additional File 2).

### Lifespan

Synchronized animals were generated by allowing gravid females/hermaphrodites (20 *C. elegans* hermaphrodites or *C. elegans fog*-*2(q71)* pseudo-females per plate; about 100 *C. inopinata* females per plate) to lay for 2-3 hours. After a few days, synchronized L4 virgin females/hermaphrodites were moved to new plates, with about 30 nematodes per plate. All animals were transferred every day for the first 4-5 days of adulthood as hermaphrodites reproduced. Subsequently, animals were scored every 1-3 days as either living or dead up until the point that all animals had died. All measurements were performed at 25°C. The number of days alive after egg-laying was taken as the measure of total lifespan. Adult lifespan was taken as the total lifespan minus two (*C. elegans*) or four (*C. inopinata*) days, as *C. inopinata* develops about twice as slowly as *C. elegans*. Statistical analyses were performed as in [36], with the *survival* package for the R statistical language being used to generate survivorship curves and the *coxme* package being used to generate Cox proportional hazard models and perform hypothesis tests (see Additional File 2).

### Fecundity

Daily offspring production was measured following overnight mating and under continuous exposure to males. For all observations, L4 *C. inopinata* NKZ2 and *C. elegans fog*-*2(q71)* animals raised at 25°C were isolated and raised for one (*C. elegans*) or two (*C. inopinata*) days to adulthood (see above). For overnight mating, single adult females/pseudo-females were shifted to the given experimental rearing temperature and mated with six males overnight. Brood sizes were measured at 15°C, 20°C, 25°C, and 30°C. The next day males were removed. Every day, embryos and larvae were counted, and egg-laying females were moved to new plates. New progeny were scored until females stopped laying for at least one (*C. elegans*) or two (*C. inopinata*) consecutive days. Continuous mating conditions were similar, except that single females were always in the presence of six males. Males that crawled up the side of the plate or otherwise died before the female stopped laying embryos were replaced with young adult males. The continuous mating observations were performed at 25°C. The instantaneous rate of natural increase [1] was calculated with Python as in [63] using life tables for *C. elegans* and *C. inopinata* constructed from the viability, fecundity, and lifespan data developed here (see Additional File 3).

### Embryo to Adult Viability

Nematode embryos were synchronized by allowing gravid females/hermaphrodites (20 *C. elegans* hermaphrodites or *C. elegans fog*-*2(q71)* pseudo-females per plate; about 100 *C. inopinata* females per plate) to lay for 2-3 hours. After the parents were removed, the number of embryos per plate were counted, and the plates were shifted to their respective rearing temperatures (15°C, 20°C, 25°C, or 30°C). L4 and adult worms were counted 4-5 days later. This fraction of mature worms/initial worm embryos was reported as the viability.

## Declarations

### Ethics approval and consent to participate

Not applicable.

### Consent for publication

Not applicable.

### Availability of data and material

All data and material not included as Additional Files are available by request.

### Authors’ contributions

GCW and PCP designed the experiments; GCW and EJ performed the experiments; GCW analyzed the results; GCW and PCP wrote the paper.

## Acknowledgements

Not applicable.

## Additional Files

Additional File 1. Supplemental Figures and Table. Figure S1. Total lifespan models with 95% confidence intervals. Figure S2. Adult lifespan models with 95% confidence intervals. Figure S3. Patterns of failed crosses across mating conditions and temperatures. Figure S4. *C. inopinata* has lower brood sizes than *C. elegans (fog*-*2)* in continuous mating conditions after removing failed crosses. Figure S5. Intersections of relevant life history trait phenotypes in *C. elegans* protein-coding genes. Table S1. Estimates of median time of developmental events.

Additional File 2. models_hypothesis_tests.R. Software for generating models and statistics.

Additional File 3. estimate_r.py. Software for estimating the rate of population increase.

Additional File 4. wormbase_phenotype_intersections.sh. Software for generating phenotype intersection UpSet plot data from a WormBase Simplemine tab-delimited file.

Additional File 5. hatch_time_data.tsv. Developmental timing data for hatching.

Additional File 6. postembryonic_milestone_time_data.tsv. Developmental timing data for postembryonic milestones.

Additional File 7. lifespan_data.tsv. Lifespan data.

Additional File 8. reproductive_duration_data.tsv. Reproductive duration data.

Additional File 9. fecundity_lifetime_access_data.tsv. Fecundity with lifetime access to males data.

Additional File 10. fecundity_overnight_mating_data.tsv. Fecundity with one overnight mating data.

Additional File 11. viability_data.tsv. Viability data.

Additional File 12. life_tables.tsv. Data used for estimating the rate of population increase.

